# Lamin B1-dependent regulation of human separase in mitosis

**DOI:** 10.1101/2024.11.28.625860

**Authors:** Julien Picotto, Francesca Cipressa, Didier Busso, Giovanni Cenci, Pascale Bertrand, Gaëlle Pennarun

## Abstract

Separase plays a central role in chromosome separation during mitosis and in centrosome cycle. Tight control of separase activity is required to prevent unscheduled resolution of sister chromatid cohesion and centrosome aberrations, thereby preserving genome stability. In mammals, despite their disassembly in early mitosis, some nuclear envelope components possess mitotic roles, but links with separase activity remain unexplored. Here, we uncover a new mechanism of separase regulation involving lamin B1, a key nuclear envelope factor. We show that separase and lamin B1 associate preferentially during early stages of mitosis. Importantly, lamin B1 depletion leads to an increase in separase recruitment on chromosomes together with premature chromatid separation, a phenotype reminiscent of separase overexpression. Conversely, similar to separase depletion, lamin B1 overexpression induces formation of diplochromosomes- resulting from chromatid separation failure-, in association with centrosome amplification. Importantly, increasing separase level prevents lamin B1-induced centrosome aberrations, suggesting a separase defect at their origin. Indeed, we show that overexpression of lamin B1 leads to a decrease in the recruitment of separase to the chromosome and a delay in its activity. Taken together, this study unveils a novel mechanism of separase regulation involving the nuclear envelope factor lamin B1, that is crucial for genome integrity maintenance.

## INTRODUCTION

In eukaryotes, mitosis marks the end of cellular division and is responsible for the fidelity of chromosome segregation in the daughter cells. Failure in this process can compromise the ploidy of the cell progeny, and be a source of tumorigenesis. The irreversible chromosome separation in anaphase is achieved by separase (ESPL1, Extra spindle pole bodies like 1 in human), a cystein protease that cleaves RAD21, a cohesin complex subunit, at centromeres (Waizenegger et al. 2002). Going through this critical stage of cell division requires a tight control of ESPL1. ESPL1 is mainly excluded from the nucleus during interphase, due in part to its nuclear export signal (NES) (Sun et al. 2006). By contrast, during prophase, it associates with mitotic chromosomes in an aurora B-dependent manner (Yuan et al. 2009), although the precise mechanism behind this recruitment remains unclear.

Despite being located on chromosomes in early mitosis, ESPL1 enzymatic activity is restricted to anaphase through its mutually exclusive binding to its inhibitory partners, securin and cyclin B1-CDK1 complex, which block its catalytic domain(Yu et al. 2021; Jallepalli et al. 2001; Stemmann et al. 2001; Hellmuth et al. 2020). During metaphase-anaphase transition, inhibition of separase activity is mainly alleviated by the degradation of securin and cyclin B1 by the ubiquitin ligase Anaphase-promoting complex/ cyclosome (APC/C), controlled by the spindle assembly checkpoint (SAC) (Yu et al. 2023; Konecna et al. 2023). A third independent inhibitory pathway regulating separase has been reported and involved its inhibitory binding to shugoshin 2-MAD2 complex, which disassembly is driven by APC/C and TRIP13/ p31comet(Hellmuth et al. 2020).

Misregulation of ESPL1 can lead to its unscheduled activation. Indeed, both early and excessive activations of ESPL1 lead to premature sister chromatid separation (PSCS), and subsequent chromosome missegregation events (Zhang et al. 2008; Shindo et al. 2022; Giménez-Abián et al. 2005; Papi et al. 2005; Hellmuth et al. 2015). Conversely, inhibition of ESPL1 results in non-disjunction of sister chromatids leading to the formation of diplochromosomes, and polyploidy (Wirth et al. 2006; Kumada et al. 2006). Furthermore, ESPL1 contributes to centrosome duplication cycle, by triggering centriole disengagement, and both its upregulation and downregulation have been associated with centrosome amplification (Galofré et al. 2020; Wirth et al. 2006). Finally, upregulation of ESPL1 expression has been described in several tumor types, including bladder, breast, bone, liver, brain or prostate cancers (Meyer et al. 2009; Mukherjee et al. 2014; Song et al. 2022; Yang et al. 2024; Zhang et al. 2008, 2024; Zhong et al. 2023). Thus, the identification of new regulators of ESPL1 activity is essential to deepen our understanding of the mechanisms involved in genomic instability associated with ESPL1 dysregulations in human cancers.

Interestingly, separase dysregulation also results in nuclear shape alterations, both in mammalian cells and in yeast (Waizenegger et al. 2002; Nakamura et al. 2002), although the link between separase and the nuclear envelope is unclear. Given that the mammalian nuclear envelope in mammals is dismantled during the early stages of mitosis, the contribution of its components to mitosis regulation have been poorly investigated. Nevertheless, several studies have reported that some nuclear envelope components have additional functions during mitosis process, including proteins of the nuclear lamina (Patil et al. 2023; Lima et al. 2024).

The nuclear lamina is a meshwork composed of lamin proteins - mainly types A/C and B-, which are type V intermediate filaments, mostly localized under the inner nuclear membranes. Mammalian mitosis is initiated by the nuclear envelope breakdown (NEB), which involves lamina disassembly by microtubule-dependent mechanical forces (Beaudouin et al. 2002; Georgatos et al. 1997) and phosphorylations of lamina proteins (Nakagawa et al. 1989; Peter et al. 1990, 1991; Heald and McKeon 1990). During the following steps of mitosis, B-type lamins contribute to spindle formation and proper chromosome segregation (Kuga et al. 2014; Tsai et al. 2006; Ma et al. 2009). In particular, depletion of lamin B1 leads to spindle assembly defects and prolonged mitosis (Tsai et al. 2006; Ma et al. 2009), but the underlying mechanisms and the contributions of lamin B1 to mitosis regulation, remain poorly understood.

Beyond its role in maintaining the structure of the nuclear envelope and in mitosis regulation, lamin B1 participates in various cellular processes, including DNA replication transcription and repair, chromatin organization, senescence, and cell migration and polarization (Dechat et al. 2008; Leeuw et al. 2018). Furthermore, we and other previously reported the involvement of lamin B1 in double-strand DNA break repair (Etourneaud et al. 2021; Butin-Israeli et al. 2015; Liu et al. 2015) and in telomere maintenance (Pennarun et al. 2021), highlighting the importance of lamin B1 to maintain genome integrity. Misregulations of lamin B1 have been reported in a wide variety of tumors, both upregulations (i.e. prostate, kidney, pancreas, liver tumors or glioma) (Luo et al. 2020; Li et al. 2013; Sun et al. 2010; Zhou et al. 2021; Gao et al. 2024) as well as down-regulations (i.e. lungs, gastric or myeloid tumors) (Jia et al. 2019; Reilly et al. 2022; Moss et al. 1999). Importantly, these dysregulations are often associated with a poor prognosis and high tumor grades. However, the underpinning mechanisms linking lamin B1 dysregulations to tumorigenesis and genetic instability remain partly elusive. Here, we report that lamin B1 is a new interacting partner of ESPL1 and that modulations of its expression alter the regulation of ESPL1 in mitosis, in association with chromosome separation defects.

## RESULTS

### Lamin B1 dysregulation interferes with chromosome separation in mitosis in human cells

Beyond nuclear envelope reassembly and disassembly, lamin B1 is involved in mitosis regulation (Tsai et al. 2006), but its precise contributions remain elusive. To characterize whether lamin B1 is involved in chromosome separation during mitosis, we first transiently overexpressed lamin B1 in normal human diploid embryonic fibroblasts (WI-38). Levels of lamin B1-overexpression achieved in WI-38 cells, as assessed by western blot (Fig. 1A), were similar to those report in diseases associated with upregulation of lamin B1 (2 to 5 fold compared to control cells), including cancers (Coradeghini et al. 2006; Li et al. 2013; Sun et al. 2010). Metaphase spread analyses performed 7-days after transfection reveal a strong significant increase in the percentage of metaphases harboring diplochromosomes - chromosomes harboring 4 chromatids joined at centromeres-in lamin B1 overexpression condition compared to the control condition (6.3% versus 0.5%, *P* < 0.005) (Fig. 1B). Of note, diplochromosomes arise after reduplication of the genome without chromatid separation, also known as endoreplication (Davoli and de Lange 2011). As this latter is often accompanied by a reduplication of centrosomes (Ganem et al. 2009; Tachibana et al. 2005; Wirth et al. 2006), we also quantified the number of centrosomes per cells by immunofluorescence imaging, using centrosome markers (centrobin or γ-tubulin). 7 days after lamin B1 overexpression in normal diploid fibroblasts, we observed a significant increase, from 3.6% to 11.9%, of cells with supernumerary centrosomes – mostly four centrosomes instead of the expected centrosome number (one or two, depending on the cell cycle phase)- (Fu et al. 2015) (Fig. 1C). The detection of centrosome reduplication in WI-38 cells is another indication that lamin B1 overexpression may induce endoreplication in normal diploid cells.

**Figure 1.**
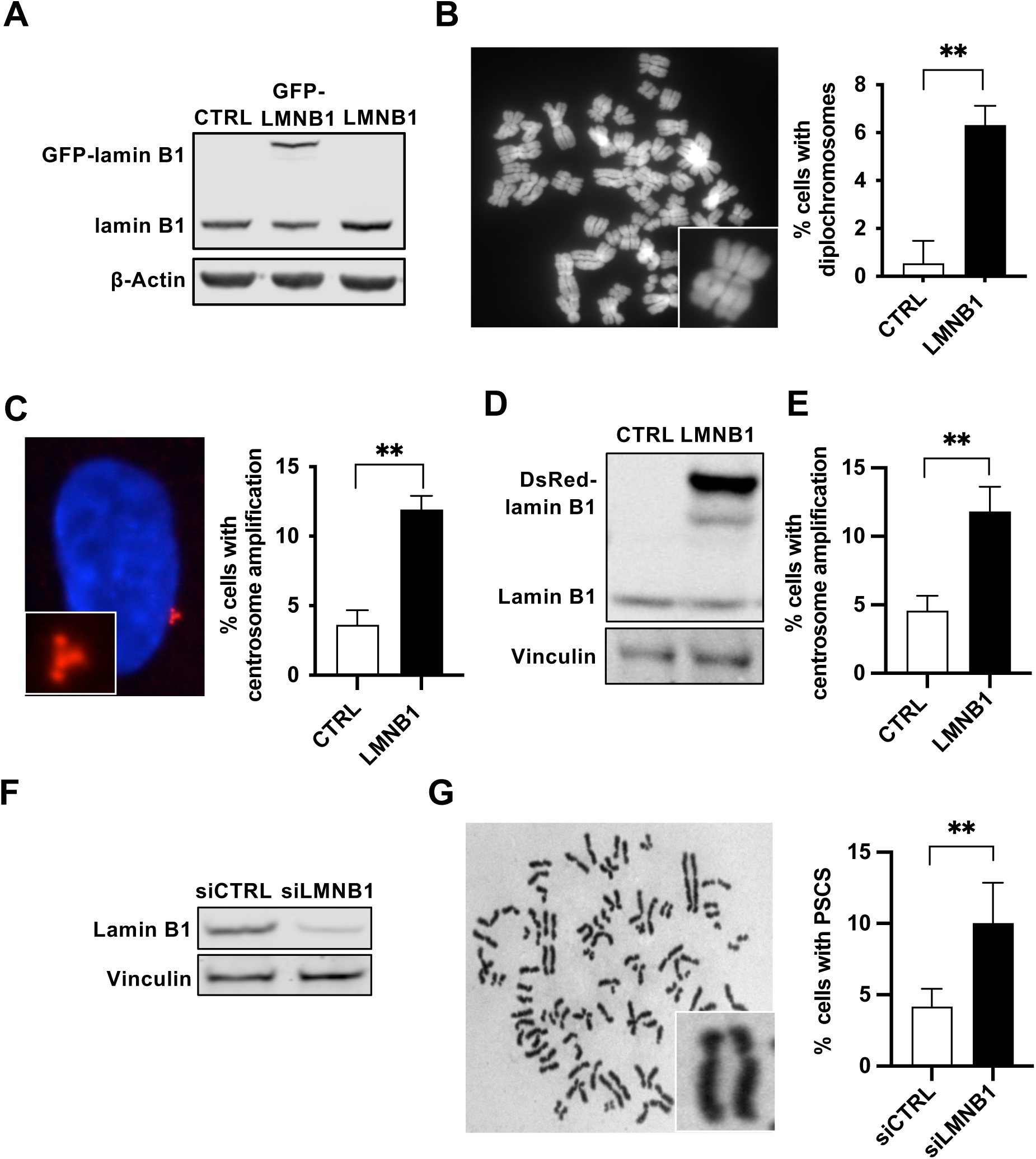
Lamin B1 dysregulation induces centrosome amplification and chromosome separation defects in human cells. (*A,B*) Induction of diplochromosomes formation after lamin B1 overexpression in human normal fibroblasts WI-38. Western blot analysis *(A)* of lamin B1 expression, 48h after transfection with an empty vector (CTRL) as control or a lamin B1 expression vector (LMNB1) in WI-38 cells or a GFP-lamin B1 expression vector (GFP- LMNB1. Representative image (*B*) of metaphase with diplochromosomes stained with DAPI from LMNB1-overexpressing cells and quantification of the percentage of cells with diplochromosomes (means ± SEM), 7 days after transfection, are shown; n= 283 and n=440 metaphases analyzed, for CTRL and LMNB1 conditions, respectively, from 3 independent experiments; ** *t*-test *P*-value = 0.0013. (*C*) Centrosome amplification induction by lamin B1 overexpression in WI-38 cells. Centrosome number per cell, 7 days after transfection as described in A, was measured by immunofluorescence using specific antibodies to centrosome (γ-tubulin). Representative images of centrosome amplification (in red and zoomed) with nuclei stained with DAPI (in blue) and quantification of centrosome amplification per cells (median ± s.d. of 3 independent experiments) are shown; \*\**t*-test *P* value = 0.0046. (*D,E*) Centrosome amplification induction after lamin B1 overexpression in human SV40- immortalized fibroblasts. Western blot analysis (*D*) of DsRed-tagged lamin B1 expression, 48h after transfection with an empty vector (CTRL) as control or a DsRed-lamin B1 expression vector (DsRed-LMNB1) in SV40-fibroblasts, using specific antibodies to lamin B1 or vinculin as loading control. Quantification (*E*) of centrosome numbers per cell (median ± s.d. of 3 independent experiments), 72h after transfection, measured by immunofluorescence using centrobin specific antibody to stain daughter centrosomes are shown; \*\**t*-test *P* < 0.005. (*F,G*) Induction of premature sister chromatid separation (PSCS) upon depletion of lamin B1 in human SV40-fibroblasts. Western-blot analysis (*F*) of lamin B1 expression, 48 h after transfection with control siRNA (siCTRL) or siRNA targeting lamin B1 (siLMNB1) in SV40- fibroblasts. Representative image (G) of a metaphase stained with Giemsa with PSCS from lamin B1-depleted cells and quantification of the percentage of cells with PSCS (means ± SEM) are shown; n > 200 metaphases, from 2 independent experiments, were analyzed for each condition; * *t*-test *P* < 0.05.

To evaluate whether this lamin B1-induced endoreplicated phenotype was not restricted to embryonic fibroblasts, we overexpressed lamin B1 in other cellular models (i.e. normal adult and transformed fibroblasts) and quantified the number of centrosomes per cells as performed for WI-38. 72h after lamin B1 overexpression, the number of cells harboring supernumerary centrosomes was significantly increased by 3-fold in transformed human fibroblasts, compared to control cells (11.8% versus 4.5%, *P* < 0.005) (Fig. 1D,E). A significant increase of extra centrosomes was also observed in adult primary fibroblasts following lamin B1 overexpression (data not shown). Together, these data indicate that lamin B1 overexpression can induce endoreplication events in both primary diploid cells and transformed cells.

To further explore the impact of lamin B1 on chromosome separation, we decreased lamin B1 level in transformed human fibroblasts by using siRNA (Fig. 1F). 72h following lamin B1 depletion, cells were analyzed for sister chromatid separation in metaphase spreads. A significant increase in metaphases with Premature Sister Chromatid Separation (PSCS) phenotype was detected in lamin B1-depleted cells compared to control cells (10% versus 4.1%, *P* < 0.05) (Fig. 1G). Thus, these data suggest that lamin B1 dysregulation impairs chromosome separation in mitosis, and that lamin B1 may be involved in the regulation of this process.

Together our results show that, while overexpression of lamin B1 could induce centrosome amplification and diplochromosomes, indicating of a potent failure in chromatid separation, its depletion could lead to premature separation of sister chromatids.

### Lamin B1 interacts with ESPL1 in mitosis

The phenotypes we observed upon lamin B1 dysregulation are reminiscent of a chromosome separation defect. Indeed, depletion of separase (ESPL1), a key protein involved in chromosome separation, leads to diplochromosomes and supernumerary centrosomes induction (Kumada et al. 2006; Wirth et al. 2006), while its overexpression or over-activation induces PSCS in mammalian cells (Meyer et al. 2009; Zhang et al. 2008). Remarkably, our observations reveal that lamin B1 dysregulation (up- and down-regulation) generates opposite phenotypes regarding chromatid separation to those caused by the dysregulation of separase. Hence, our results prompt us to hypothesize that lamin B1 might be a new regulator of ESPL1 in mitosis. In order to investigate this hypothesis, we first evaluated the ability of lamin B1 and ESPL1 to interact in human cells. Immunoprecipitation using specific antibody against ESPL1 revealed that endogenous lamin B1 co-precipitates with endogenous ESPL1 (Fig. 2A, *upper panel*). Conversely, endogenous ESPL1 co-precipitates with endogenous lamin B1 in immunoprecipitation using an antibody against this later (Fig. 2A, *lower panel*). Of note, as both ESPL1 and lamin B1 have properties to bind DNA (Sun et al. 2009; Pascual-Reguant et al. 2018), co-immunoprecipitation experiments were performed in presence of benzonase to exclude potent protein interactions *via* DNA bridges.

**Figure 2.**
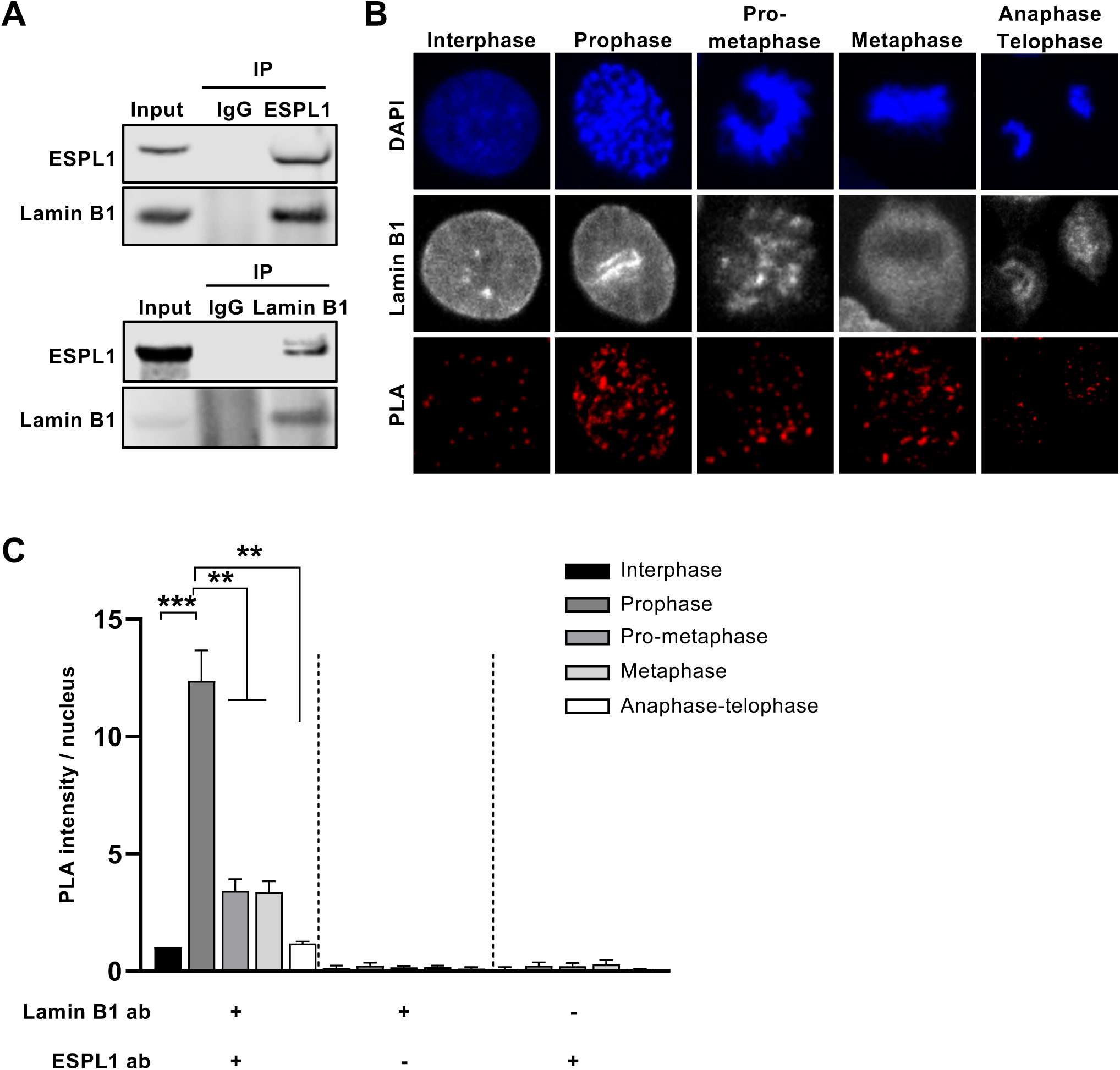
Endogenous lamin B1 and ESPL1 interact preferentially in early mitosis. (*A*) Co-immunoprecipitation of endogenous ESPL1 and lamin B1 in human cells. Lysates of non-transfected SV40-human fibroblasts pretreated with benzonase nuclease were immunoprecipitated with anti-ESPL1 (*upper panel*) or anti-lamin B1 antibodies (*lower panel*) or IgG as controls. Immunoprecipitates were revealed by western blotting using antibodies specific to ESPL1 and lamin B1 antibodies. Similar results were obtained in three independent experiments. (*B,C*) *In situ* interaction between ESPL1 and lamin B1 during cell cycle. *In situ* proximity ligation assay (PLA) as a function of cell cycle was performed on non-transfected human SV40-fibroblasts using specific antibodies against lamin B1 and ESPL1 and DAPI staining to sort cells into five stages of the cell-cycle: interphase, prophase, pro-metaphase, metaphase and anaphase-telophase. Representative images (*B*) of PLA (in red), lamin B1 (in grey) and DAPI staining (in blue) of cells and quantification (*C*) of ESPL1-lamin B1 PLA intensity per nuclei (median ± s.d.) in the five categories from 3 independent experiments are shown. PLA negative controls with only one of the antibodies against ESPL1 or lamin B1 are shown (ESPL1 ab or LB1 ab); n>30 cells per condition; \*\**t*-test *P* values < 0.01; *** P < 0.001.

To study the cellular localization and dynamics of this new identified interaction between ESPL1 and lamin B1, we performed Proximity Ligation Assay (PLA). We first observed *in situ* close proximity between endogenous lamin B1 and ESPL1 in interphase nuclei (Fig. 2B,C). Given that ESPL1 associates with chromosomes from prophase to anaphase (Yuan et al. 2009), and that its catalytic activity is mainly restricted to anaphase (Kamenz and Hauf 2017), we hypothesized that lamin B1 and ESPL1 associate in a cell-cycle dependent manner. Therefore, we classified the cells according to five cell cycle phases: interphase, prophase, pro-metaphase, metaphase, or anaphase/telophase. This analysis revealed that the close proximity (PLA intensity signal) between ESPL1 and lamin B1 is significantly higher in prophase nuclei (12-fold increase) compared to interphasic nuclei, then decreases during pro-metaphase and metaphase (3.5-fold increase compared to interphase cells), and finally returns to a level comparable to that of interphase cells (Fig. 2C).

To evaluate the effect of high lamin B1 expression on its association with ESPL1, we performed PLA experiment after lamin B1 overexpression in transformed human fibroblasts (Fig. 3). These experiments revealed that the level of association between ESPL1 and lamin B1 significantly increases in cell overexpressing lamin B1, with a positive correlation according to the expression level of lamin B1. The intensity of ESPL1-lamin B1 PLA was 2.5-fold higher in cells with 2 to 5-fold lamin B1 overexpression (Fig. 3B). We next investigated the dynamic of this interaction upon lamin B1 overexpression during cell cycle. We found that that lamin B1 overexpression induces an enhanced interaction with ESPL1, preferentially in prophase (Fig. 3C). These data indicate that lamin B1 preferentially associates with ESPL1 during mitosis, especially in prophase, and to an even higher extent upon lamin B1 overexpression. Altogether our data unveil a new interaction between lamin B1 and ESPL1 and suggest that lamin B1 may be a regulator of ESPL1 in mitosis.

**Figure 3.**
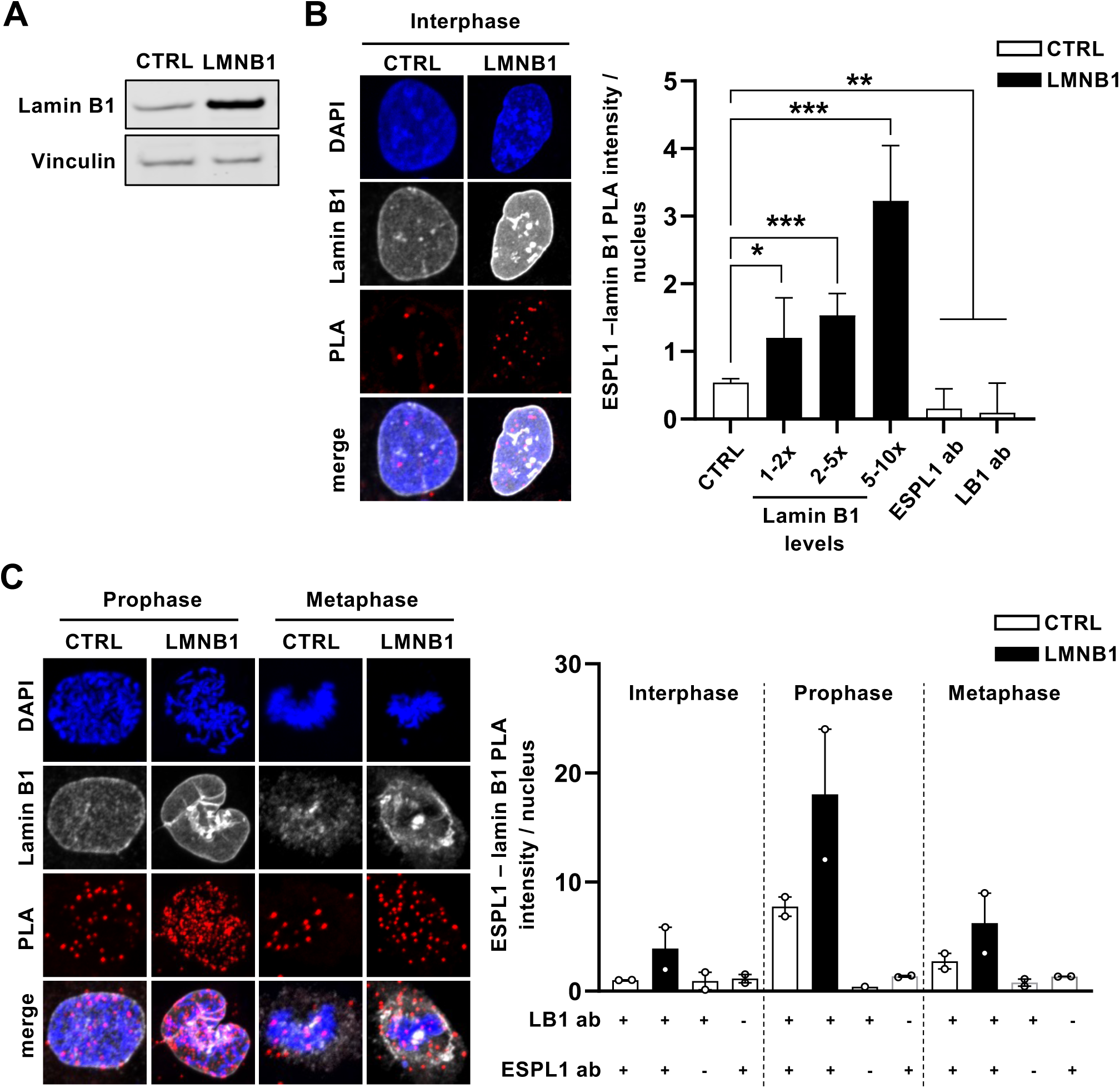
Increase of laminB1-ESPL1 interaction upon lamin B1 overexpression. (*A,B*) *In situ* interaction between lamin B1 and ESPL1 upon lamin B1 overexpression. Lamin B1 protein level (*A*) analyzed by western blot in human SV40-fibroblasts, 48h after transfection with an empty vector (CTRL) as control or a lamin B1 expression vector (LMNB1) is shown. (*B*) ESPL1-Lamin B1 *in situ* proximity assay (PLA), as a function of lamin B1 intensity, coupled with lamin B1 immunofluorescent staining was performed in transfected cells as described in (*A*). Lamin B1-overexpressing cells were classified in three categories according to their level of lamin B1: 1–2×, 2–5×, 5–10×, compared to that of control cells. Representative images of ESPL1-lamin B1 PLA (in red) and quantification of PLA dots per nuclei (median ± s.d.) from 3 independent experiments are shown. PLA negative controls with only one of the antibodies against ESPL1 or lamin B1 (ESPL1 ab or LB1 ab) are shown; n>50 cells; **t-test *P* value < 0.01; *** *P* < 0.001. (*C*) Impact of lamin B1 overexpression on *in situ* ESPL1-Lamin B1 interaction during cell cycle. PLA assay performed in transfected cells as described in (*A*) was analyzed as a function of cell cycle phases: DAPI staining of control (CTRL) or lamin B1 (LMNB) overexpressing human SV40-fibroblasts was used to sort cells into three categories of cell-cycle phases: interphase, prophase, post-prophase (prometaphase, metaphase, and anaphase-telophase transition). Representative images of PLA staining and quantification of ESPL1-lamin B1 PLA intensity per nuclei in the three categories from 3 independent experiments is shown (median ± s.d.). PLA negative controls with only one of the antibodies against ESPL1 or lamin B1 are shown (ESPL1 ab or LB1 ab).

### Lamin B1 dysregulation modifies ESPL1 localization in mitosis

Considering that lamin B1 is a new binding partner of ESPL1 and that its overexpression increases this association, we hypothesized that lamin B1 dysregulation might alter ESPL1 regulation in mitosis. Hence, we first analyzed the impact of lamin B1 inhibition on the intracellular localization of ESPL1 protein during mitosis by immunofluorescence imaging in transformed fibroblasts (Fig. 4B). Immunofluorescence imaging of ESPL1 revealed that lamin B1-depleted cells in mitosis exhibit a significant increase of ESPL1 signal on chromosomes (1.7-fold, compared to control cells, 48h following lamin B1 depletion), whereas no impact was observed in interphase nuclei (Fig. 4B). To analyze whether this effect is due to an increase of ESPL1 protein expression or to an alteration of ESPL1 localization, we measured the level of ESPL1 protein by western blotting in synchronized cells by a double thymidine block. This revealed that depletion of lamin B1 did not alter ESPL1 expression level, neither in asynchronous cells nor in mitotic cells (Fig. 4C,D). Of note, lamin B1 depletion had no significant effect on cell synchronization in mitosis, assessed by the quantification of the phosphorylation of histone H3 on serine 10 (pH3^ser10^) (Fig. 4E). Altogether, these data indicate that lamin B1 depletion leads to an alteration of ESPL1 localization by inducing an enrichment of ESPL1 on chromosomes during mitosis.

**Figure 4.**
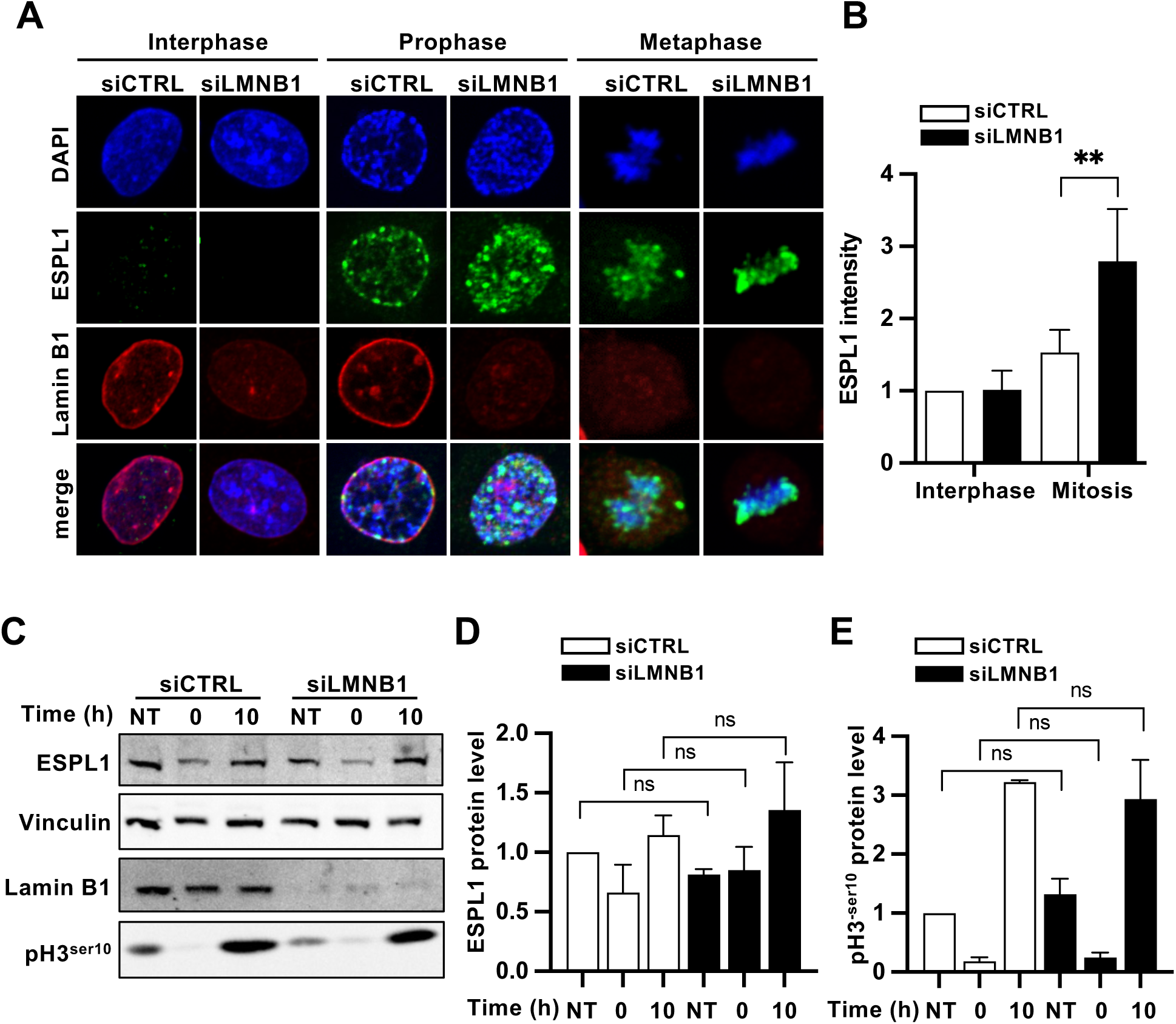
Lamin B1 depletion increases ESPL1 localization on chromosomes during mitosis. (*A,B*) Increase of ESPL1 intensity in mitosis upon lamin B1 depletion. ESPL1 intensity as a function of cell cycle phases was analyzed by immunofluorescence in human SV40- fibroblasts, 48 h after transfection with siRNA against lamin B1 (siLMNB1) or control siRNA (siCTRL). DAPI staining of human SV40-fibroblasts was used to sort cells mitosis or in interphase. *R*epresentative images (*A*) of immunostaining of ESPL1 (in green) and lamin B1 (in red) and nuclei stained with DAPI (in blue) and quantification (*B*) of ESPL1 intensity per cell in interphase or mitosis calculated by normalizing to ESPL1 intensity of control siRNA cells in interphase (set as 1) are shown; n>30 cells, median ± s.d. of 3 independent experiments; \*\**t-* tests *P* value < 0.01; *** *P* < 0.001. (*C-E*) Impact of lamin B1 depletion on ESPL1 protein level in SV40-fibroblasts synchronized in S-phase and mitosis. Cells transfected as described in (*A*) and synchronized by a double thymidine block were harvested just after the block (0) – cells synchronized in S phase - or 10 h after (10) – mitotic cells- and subjected to western blot analysis (*C*) with ESPL1 and lamin B1 antibodies; anti-phospho-H3 Ser10 (pH3^ser10^) antibody was used as a mitotic control and vinculin as loading control. Quantifications of ESPL1 (*D*) and phospho-H3 Ser10 (*E*) protein levels from 3 independent experiments (median ± s.e.m.) are shown; ns= *t*-test *P* value non-significant.

To further characterize the impact of lamin B1 dysregulation on separase regulation, we evaluated whether lamin B1 overexpression may have a reverse impact on ESPL1 localization, meaning a decrease of ESPL1 presence on chromosomes. Immunofluorescence imaging revealed that ESPL1 signal on mitotic chromosomes decreases by 30%, 48h following lamin B1 overexpression, compared to control cells (Fig. 5A,B). To exclude that this decrease results from an alteration of in ESPL1 protein level, we measured its level by western blotting. This experiment revealed that lamin B1 overexpression did not affect ESPL1 protein level throughout the cell cycle (Fig. 5C-E), suggesting that the observed decrease of ESPL1 signal on chromosomes is rather due to an alteration of its localization upon lamin B1 depletion.

**Figure 5.**
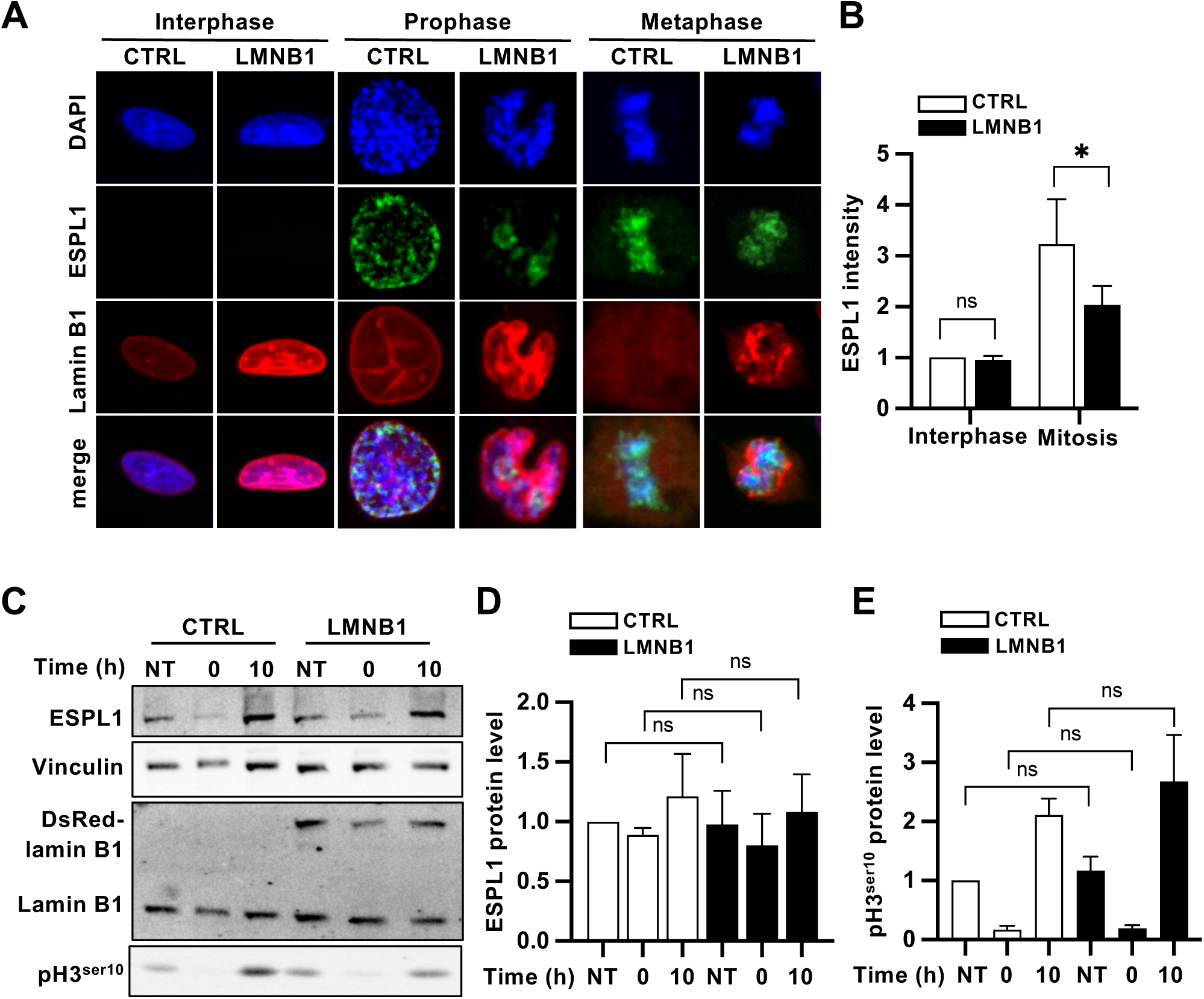
Lamin B1 overexpression decreases ESPL1 presence on chromosomes during mitosis. (*A*,*B*) Decrease of ESPL1 intensity in mitosis upon lamin B1 depletion. ESPL1 intensity as a function of cell cycle phases analyzed by immunofluorescence in human SV40- fibroblasts, 48 h after transfection with CTRL, LMNB1 or Dsred-LMNB1 expression vectors. DAPI staining of human SV40-fibroblasts was used to sort cells in mitosis or in interphase. Representative images (*A*) of immunostaining of ESPL1 (in green) and lamin B1 (in red) and nuclei stained with DAPI (in blue) and quantification (*B*) of the relative ESPL1 intensity for each category calculated by normalizing to ESPL1 intensity of control vector cells in interphase; n=6 independent experiments and n>30 cells analyzed per condition; bars show median ± s.e.m.; \**t*-test *P* < 0.05. (*C-E*) Impact of lamin B1 overexpression on ESPL1 protein level human SV40-fibroblasts synchronized in S-phase and mitosis. Cells transfected as described in (*A*) and synchronized by a double thymidine block were harvested just after the block (0) – cells synchronized in S phase - or 10 h after (10) – mitotic cells- and subjected to western blot analysis with ESPL1 and lamin B1 antibodies; anti-phospho-H3 Ser10 (pH3^ser10^) antibody was used as a mitotic control and vinculin as loading control. Quantifications of ESPL1 (*D*) and phospho-H3 ser10 (*E*) protein levels from 4 independent experiments (median ± s.d.) is shown; ns= *t*-test *P* value non-significant.

These alterations of ESPL1 localization during mitosis upon lamin B1 dysregulation suggest that lamin B1 could play a role in the timely regulation of ESPL1 association with chromosomes, and subsequent activity in anaphase.

### ESPL1 mitotic activity is modified after lamin B1 modulations

To characterize whether the differences in the abnormal localization of ESPL1 in mitosis following lamin B1 dysregulation results in alterations of separase activity, we used HeLa- cells constitutively expressing a fluorescent substrate (HeLa-sensor, kindly provided by Dr. Olaf Stemmann (Henschke et al. 2019)), which allows the measurement of ESPL1 enzymatic activity in time-lapse fluorescent microscopy. This substrate (designed by Turo Hirota (Shindo et al. 2012) consists of a RAD21 region containing its ESPL1 cleavage sites, fused in C-terminal to a GFP tag and in N-terminal to a mCherry tag, followed by a H2B fusion protein to locate the construct along the chromosome arms (Fig. 6A). The activation of ESPL1 leads to the cleavage of this construct, and in the loss of the GFP tag while the mCherry remains on the chromosomes through the H2B fusion protein (Shindo et al. 2012).

**Figure 6.**
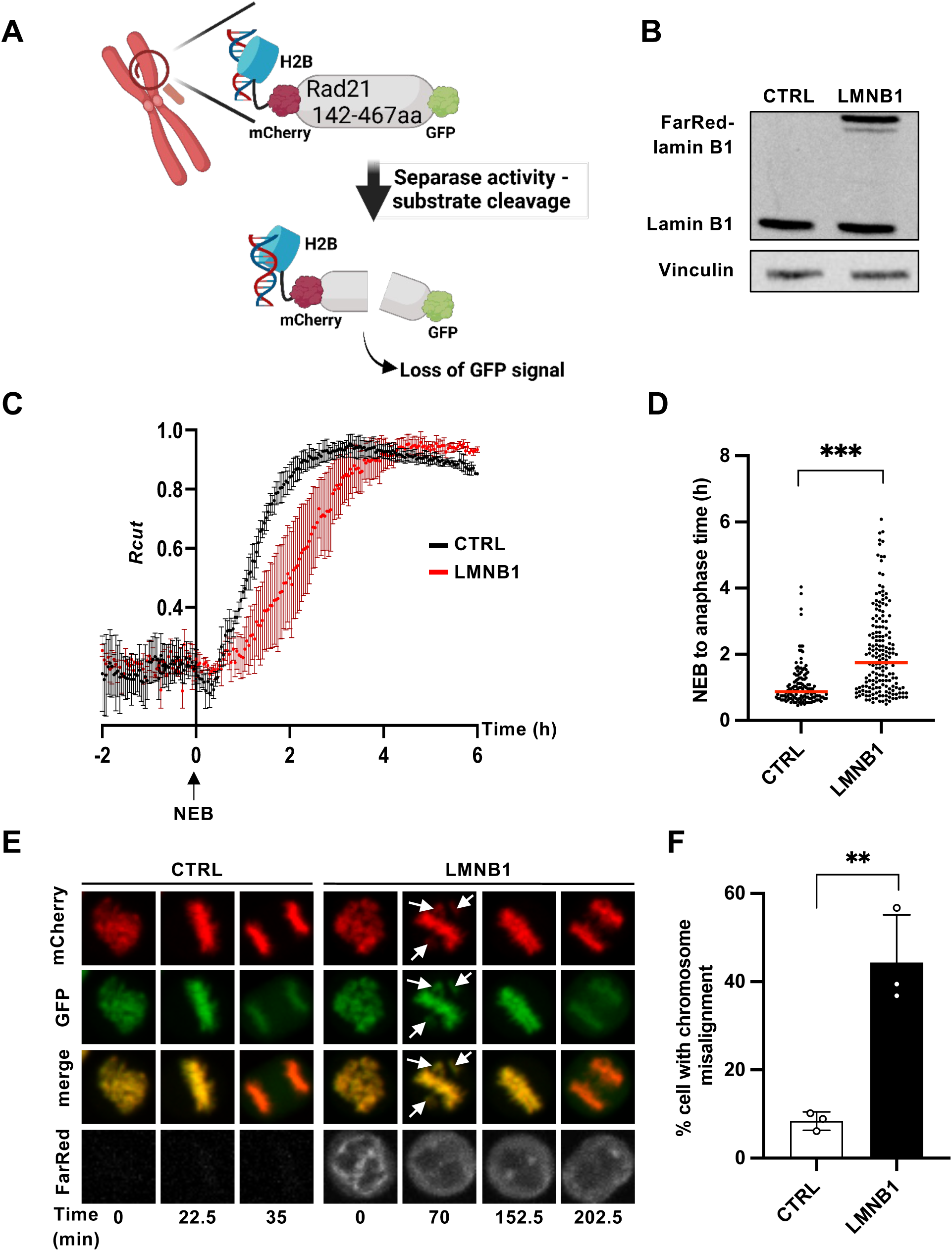
Lamin B1 overexpression alters ESPL1 activity in mitosis. (*A*) Scheme of the separase substrate (described by Dr. Toru Hirota, in Shindo et al. 2012) to measure Separase activity *in cellulo*: cleavable Rad21 fragment fused with mCherry tag and H2B in N-ter, and with GFP tag in C-ter. This scheme was created with BioRender.com. (*B*) Overexpression of lamin B1 in HeLa cells stably expressing the separase substrate described in (*A*). Western blot analysis of lamin B1 expression levels, 48 h after transfection with FarRed-LMNB1 (LMNB1) or CTRL vectors; representative image is shown; similar results were obtained in 3 independent experiments. (*C*) Separase activity followed by time-lapse upon lamin B1 overexpression in HeLa cells. Synchronized cells, transfected as described in (*B*) were imaged by time-lapse microscopy at the onset of anaphase. The graph represents cumulative normalized Rcut values (median ± s.d), reflecting separase activity, from lamin B1-overexpressing cells (LMNB1) and control cells (CTRL) during mitosis from 3 independent experiments; n > 35 mitosis analyzed per each condition per each experiment. The Nuclear Envelope Breakdown (NEB) onset was considered as time 0 h. (*D*) Delay between nuclear envelope breakdown and anaphase onset after lamin B1 overexpression. Quantification of the time between the NEB and anaphase onset in cells described in (*C*). Graph represents the medians ± s.d. of 3 independent experiments; *** *t*-tests *P* value <0.0001. (*E*) Representative images of the time-lapse imaging described in (*C*); white arrows showing misaligned chromosomes. (*F*) Quantification of the percentage of mitosis with chromosome misalignment (medians ± s.d) after lamin B1 overexpression. Chromosome misalignment was determined for each cell analyzed in (*C*) as persistent unattachment of chromosomes in anaphase (showed by white arrows in (*E*); n= 3 independent experiments, ** *t-*test *P* < 0.005.

Hence, we performed time-lapse fluorescent microscopy in synchronized HeLa-sensor cells, 48h after overexpression of FarRed-tagged lamin B1 (Fig. 6B). We analyzed separase activity in tracked mitosis by measuring the Rcut values as described by Shindo et al. (2012), which provides a precise measure of the cumulative substrate cleavage events, reflecting ESPL1 activity. The analysis revealed a significant delay of ESPL1 activation in lamin B1- overexpressing cells (Fig. 6C). In addition, we observed an extension of the time between the NEB and the onset of anaphase, from a mean of 45 min in control cells to almost 2 h in lamin B1-overexpressing cells (Fig. 6D). Moreover, these effects were associated with a significant 5.2- fold increase in chromosome misalignment (44.3% in lamin B1-overexpressing cells compared to 8.4% in control cells) (Fig. 6E,F), suggesting that lamin B1 overexpression alters separase activity in mitosis, leading to an unscheduled chromosome separation.

Together, these findings indicate that lamin B1 overexpression leads to an increase in its interaction with ESPL1, in association with an alteration of the localization and activity of this latter. We therefore hypothesized that lamin B1 in excess could result in ESPL1 trapping, thereby impairing its functions in anaphase.

### ESPL1 defect causes centrosome amplification upon lamin B1 overexpression

Disruption of ESPL1 activity has been reported to result in failure of sister chromatid separation and centrosome amplification (Wirth et al. 2006; Kumada et al. 2006). As lamin B1 overexpression induces chromosome separation defects and centrosome amplification, we next investigated whether lamin B1-induced centrosome amplification could arise from a separase defect. To achieve this purpose, we performed complementation experiments by overexpressing ESPL1 in addition to lamin B1 in order to test whether elevated level of ESPL1 could prevent lamin B1-induced centrosome amplification to occur. We overexpressed lamin B1 and ESPL1 in transformed human fibroblasts and quantified the rate of centrosome amplification by immunofluorescence imaging (Fig. 7). Overexpression of lamin B1 and ESPL1 were assessed by both western blotting (Fig. 7A) and immunofluorescence quantification of their DsRed or HA tag, respectively (Fig. 7,C). Further analysis were performed in cell population with comparable levels of lamin B1 overexpression. 72h after transfection, we observed that the rate of centrosome amplification was significantly reduced by 2-fold in cells overexpressing both lamin B1 and ESPL1, compared to cells overexpressing only lamin B1 (11.9 % ± 1.9 versus 5.7% ± 2.8%) and recovers a similar level to that of control cells (Fig. 7D). It is noteworthy that double-transfected cells did not seem to show striking cell cycle defects, and displayed a similar distribution of cells with 1 or 2 centrosomes compared to control cells (Fig. 7E). These data suggest that lamin B1 overexpression-induced centrosome amplification could result from a separase defect. Altogether, our data indicate that lamin B1 overexpression by altering separase localization and activity, lead to a defect in separase, thereby leading to chromatid separation defect and centrosome amplification.

**Figure 7.**
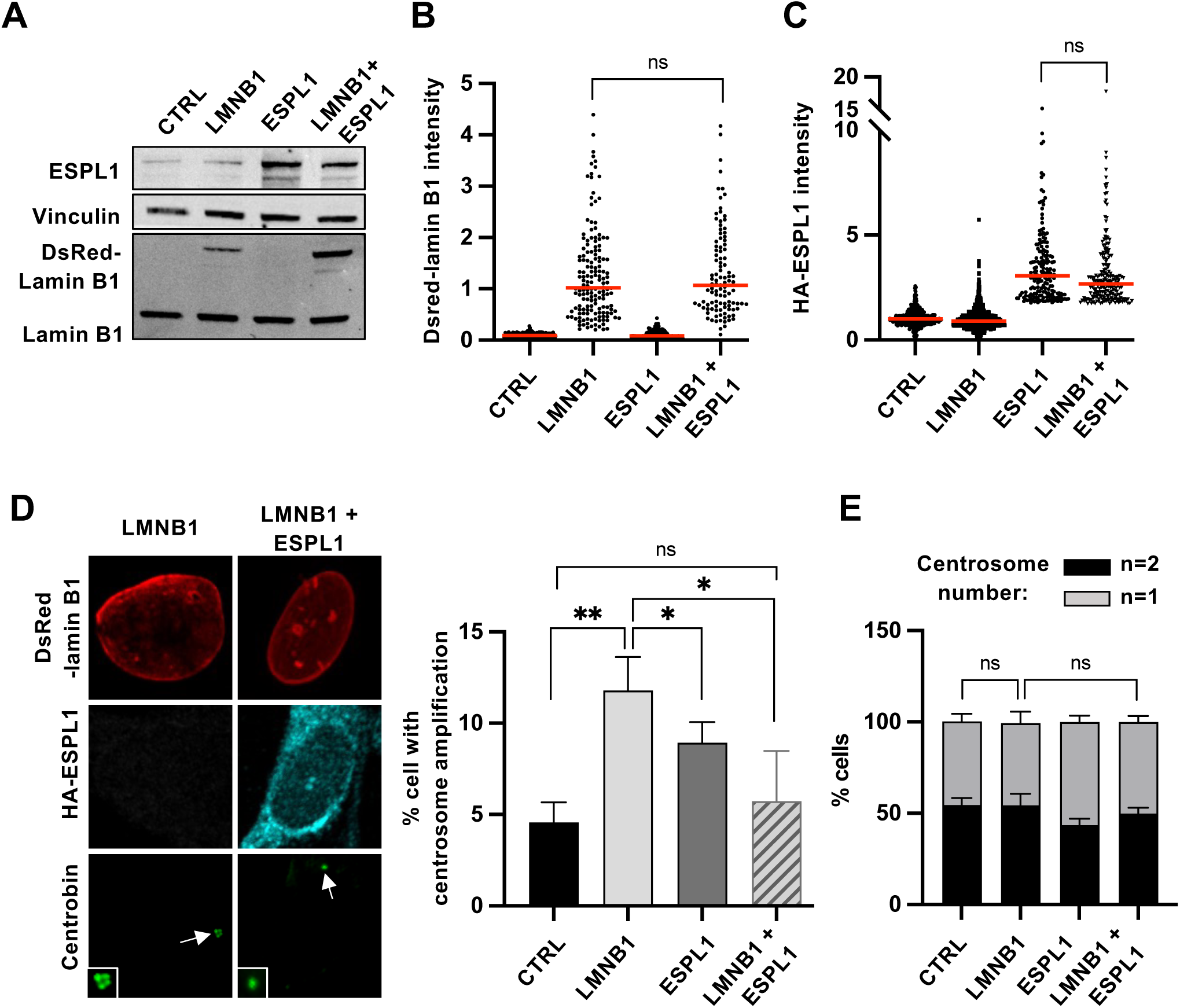
Ectopic expression of ESPL1 prevents lamin B1–induced centrosome amplification. (*A*) SV40-fibroblasts were transfected either with Dsred-LMNB1 (LMNB1) or HA-ESPL1 (ESPL1) or CTRL vectors or co-transfected with both HA-ESPL1 and DsRed-LMNB1 (LMNB1 + ESPL1) vectors. 48 h after transfections, HA-ESPL1 and DsRed-lamin B1 expression levels were analyzed by western blots using specific antibodies against ESPL1, lamin B1 and vinculin as loading control. n=3 independent experiments. (*B*) Quantification of the expression level of DsRed-lamin B1 by fluorescent imaging of Dsred signal in cells further analyzed for centrosome number in (*D*); medians in red of 3 independent experiments; note that the levels of Dsred in LMNB1 and LMNB1 + ESPL1 conditions are similar: *t-*test *P* value non-significant (ns). (*C*) quantification of the expression level of HA-ESPL1 by fluorescent imaging of HA-tag signal in cells further analyzed for centrosome number in (*D*); medians in red of 3 independent experiments; note that the levels of HA-tag in ESPL1 and LMNB1 + ESPL1 conditions are similar: *t-*tests *P* value = ns. (*D*) Centrosome amplification in rescue experiments described in (*A*). Centrosome number per cell, 72h after transfection, was measured by immunofluorescence using an antibody to centrobin to stain daughter centrosomes. Cells expressing only HA-ESPL1, only DsRed-LMNB1 or both proteins were analyzed and compared to control condition (CTRL). *Left panel,* representative staining of centrosomes (in green and pointed by arrows) in cells expressing DsRed-lamin B1 (in red) (LMNB1) or both HA-ESPL1 (in cyan) and DsRed-lamin B1 (LMNB1 = ESPL1). *Middle panel*, quantification of cells with centrosome amplification (3 or 4 centrosomes) is shown (n>30 cells, median ± s.e.m. of 3 independent experiments); ** t-tests *P* value < 0.01; *** *P* < 0.001. (*E*) Distribution of cells harboring 1 or 2 centrosomes among the rescue experiment conditions as described in (*A*); note that cells with centrosome amplification are not shown here; median ± s.d. of 3 independent experiments are shown; ns= t*-*test *P* value non-significant.

## DISCUSSION

Until now, the relationship between separase and the nuclear envelope has been poorly explored. However, previous studies have shown that separase silencing in human cells, or overexpression of a C-terminal fragment of separase, induces nuclear shape alteration (Nakamura et al. 2002; Waizenegger et al. 2002), but the underlying mechanisms of this unexpected link remain elusive. Despite that the nuclear envelope breaks down in mitosis, several nuclear envelope components, beyond theirs role in nucleus architecture, have been reported to play a role in mitosis regulation, including B-type lamins, which contribute to spindle assembly and chromosome segregation (Tsai et al. 2006; Ma et al. 2009; Kuga et al. 2014). But whether nuclear envelope components are involved in separase regulation during mitosis, or how nuclear envelope defects could impact separase activity, has not been investigated yet.

Here we report a novel molecular link between a nuclear envelope protein, lamin B1, and separase (ESPL1) in mammals. Indeed, we found that lamin B1 and ESPL1 interact in human cells. No evidence of interaction between other mammalian nuclear envelope components and separase have been reported so far. Supporting our findings, *Drosophila* separase SSE interacts with lamin Dm0, the homologue of B-type lamin in this fly (Cipressa F. and Cenci G. in preparation), showing that this uncovered interaction between lamin and separase are conserved through evolution. We further show that lamin B1 and ESPL1 preferentially associate during early mitosis, and that lamin B1 overexpression increases this association, suggesting that lamin B1 modulations may impact separase functions. Supporting this idea, we found that lamin B1 dysregulation mirrors the chromosome segregation phenotypes induced upon separase dysfunction. Indeed, lamin B1 depletion induces PSCS, a phenotype reminiscent of separase overexpression (Zhang et al. 2008). By contrast, as observed upon separase inhibition, lamin B1 overexpression leads to an endoreplication phenotype, as evidenced by diplochromosomes and supernumerary centrosomes, indicating that lamin B1 dysregulation may impair chromosome segregation by altering separase regulation (Wirth et al. 2006; Kumada et al. 2006).

The nuclear envelope, especially the inner nuclear membrane and the lamina, has been proposed to play a role in the regulation of nucleoplasmic proteins through their sequestration (Serebryannyy and Misteli 2018). In particular, lamin B1 overexpression has been reported to induce the sequestration of several of its interactors, such as transcription factors, DNA repair factors or telomeric proteins (Pennarun et al. 2021; Etourneaud et al. 2021; Malhas et al. 2009), resulting in an impairment of their functions. As we showed that lamin B1 overexpression leads to an increased interaction with ESPL1 during mitosis, we propose that lamin B1 in excess could sequester separase and disturb its regulation in mitosis, resulting in defects in sister chromatids disjunction, and centrosome amplification (Fig. 8). Conversely, lamin B1 depletion might induce a lack of ESPL1-lamin B1 interaction as well as a misregulation of ESPL1 recruitment to mitotic chromosomes, that might, in turn, cause a deficiency in ESPL1 regulation between prophase and anaphase, which could result in its untimely activation, and PSCS phenotype.

**Figure 8.**
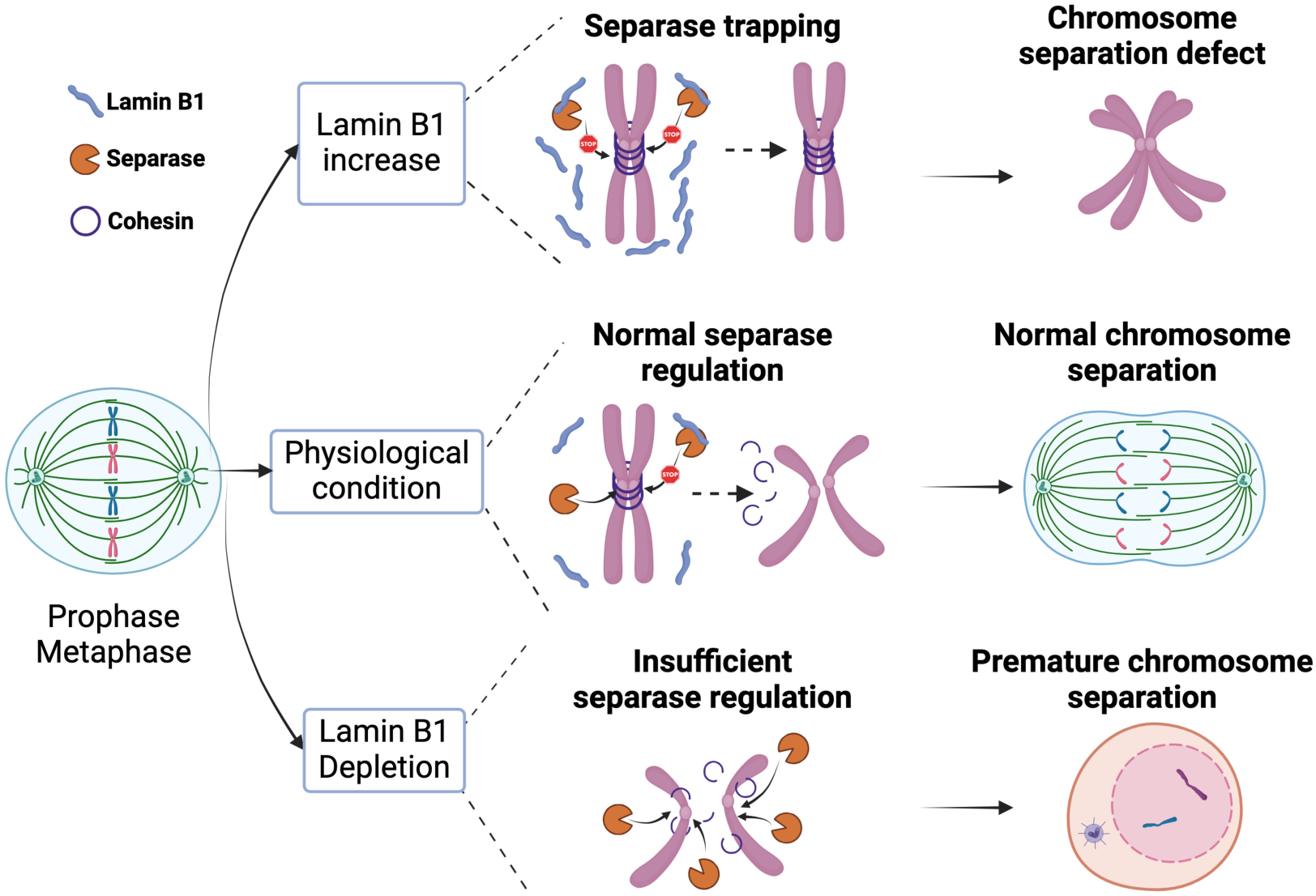
Model for the regulation of ESPL1 by lamin B1 during mitosis. In wild-type cells, we propose that lamin B1 interacts with ESPL1 to regulate its activity before anaphase to ensure a proper and timely disjunction of sister chromatids. Upon lamin B1 overexpression, the association between lamin B1 and ESPL1 is strongly increased during mitosis. ESPL1 could therefore be trapped by the excess of lamin B1, leading to the decrease of its recruitment on mitotic chromosomes, which could, in turn, results in chromosome separation defects (i.e. non-disjunction of chromosomes-followed by diplochromosomes at the next mitosis-, lagging chromosomes, chromosomes misalignments, supernumerary centrosomes). By contrast, lamin B1 depletion induces an increase in the recruitment of ESPL1 on chromosomes in early mitosis, which could induce the premature separation on sister chromatids. This model was created with BioRender.com.

Besides the previously described mechanisms of separase regulation, involving post-transcriptional modifications and inhibitory interactions with securin, cyclin B1 and SGO2 (Waizenegger et al. 2002; Holland and Taylor 2006; Hellmuth et al. 2020), we propose that lamin B1, through its interaction with ESPL1 in early mitosis, may constitute a novel layer of separase regulation that contributes to well orchestrate chromatid separation in mitosis. Supporting this model, we showed by using a specific fluorescent substrate of separase activity, that lamin B1 in excess, in addition to enhance its interaction with separase, and alter the localization of this latter on mitotic chromosomes, delays separase activation and anaphase onset. Prolonged arrest in metaphase has been previously reported to lead to an exit of mitosis without cytokinesis (Davoli and de Lange 2011), that can be a source of endoreplication. In accordance, we observed a phenotype of endoreplication, marked by diplochromosomes formation and supernumerary centrosomes, following lamin B1 overexpression, suggesting that this phenotype could be linked to a defect in separase activity. Importantly, lamin B1-induced supernumerary centrosomes are prevented by increasing the protein level of separase. Therefore, collectively, these data suggest that lamin B1 dysregulation may specifically impact separase activity, leading to chromatid separation failure, and subsequent mitotic slippage, centrosome amplification and tetraploid cells (Ganem et al. 2009).

The nuclear envelope breakdown at the initiation of mitosis and its reformation during mitosis exit, has for long been considered as the only involvements of lamins in mitosis. However, several studies revealed that B-type lamins facilitate mitosis by assembling as a matrix, under the regulation of Nudel and dynein proteins (Tsai et al. 2006; Ma et al. 2009). Moreover, depletion of lamin B2 in human cells has been shown to induce aneuploidy, mitotic defects and alters spindle formation (Kuga et al. 2014), but the underlying mechanisms are still unknown. Here, we report a new implication of lamin B1 in mitosis through its interaction with ESPL1, showing a novel role for lamin B1 during mitosis, crucial for chromosome segregation beyond the nuclear breakdown and spindle assembly.

Furthermore, the regulation of ESPL1 recruitment to mitotic chromosomes is still partially understood. Indeed, ESPL1 is excluded from the nuclei during interphase and detected in association with chromosomes during prophase in a Aurora B-dependent manner (Yuan et al. 2009), even if no phosphorylation site has been identified. Since lamin B1 dysregulation alters separase intensity on chromosomes during mitosis, we could speculate that the interaction we detected in endogenous conditions between the two proteins contributes to the regulation of separase association with chromosomes. However, whether lamin B1 dysregulation could also impact the other pathways of separase regulation involving its inhibitory partners remains to be investigated.

We show that elevation of separase level can rescue the induction of supernumerary centrosomes in transformed fibroblast upon lamin B1 overexpression, indicating that a separase defect is involved in this phenotype. We and others have shown the lamin B1 overexpression also alters DNA repair mechanisms and telomere protection (Pennarun et al. 2021; Etourneaud et al. 2021; Butin-Israeli et al. 2015; Liu et al. 2015). These phenotypes can also induce tetraploidization, lagging chromosomes or chromosome misalignment during metaphase (Reina-San-Martin et al. 2005; Wilsker et al. 2012; Yoshihara et al. 2004; Hockemeyer et al. 2006; Kibe et al. 2010; Lazzerini Denchi et al. 2006). Hence, we cannot exclude that DNA repair defects and telomere alterations may contribute to the phenotypes we observed in this study. It is also important to consider that these mechanisms could coexist in cells and overlap in some contexts. Indeed, the extension of mitosis can be both a cause and a consequence of ESPL1 dysregulation, associated with lagging chromosomes and chromosome misalignment (Shindo et al. 2022; Giménez-Abián et al. 2005; Papi et al. 2005).

It has been previously described that centrosome amplification is sufficient to promote tumorigenesis (Godinho et al. 2014; Basto et al. 2008; Levine et al. 2017). However, the tetraploid state can be transient in cancer, and excessive centrosomes can be lost along successive cell cycles (Galofré et al. 2020; Zhong et al. 2023). In the other hand, numerous studies have reported that cells can arrest in G1, enter senescence or trigger apoptosis after tetraploidization (Wirth et al. 2006; Andreassen et al. 2001; Storchova and Kuffer 2008; Panopoulos et al. 2014; Castedo et al. 2006; De Santis Puzzonia et al. 2015). Nonetheless, in some contexts, primary cells can become tetraploid or aneuploid after successive division cycles, despite the presence of p53 (Potapova et al. 2016; Kuznetsova et al. 2015). Here, we observed centrosome amplification after lamin B1 overexpression in both transformed and primary cells, suggesting that this effect might be independent of p53.

Lamin B1 overexpression, correlated with poor prognosis and tumor grade, has been observed in many cancer types (Evangelisti et al. 2022; Irianto et al. 2016), while the implications of this overexpression on tumorigenesis remain elusive. In addition to telomeric damages and DNA repair defects (Etourneaud et al. 2021; Pennarun et al. 2021; Butin-Israeli et al. 2015), we now show that lamin B1 dysregulation results in supernumerary centrosomes and mitotic defects (diplochromosomes and chromosome misalignment), and could have dramatic effect on ploidy. These phenotypes are important sources of genomic instability and could contribute to tumorigenesis, and could thus explain why lamin B1 dysregulation in cancer is often correlated with poor prognosis and high tumor grade.

In conclusion, our results uncover a novel molecular link between lamin B1 and ESPL1, and show the impact of lamin B1 dysregulation on ESPL1 activity, highlighting a novel implication of lamin B1 during mitosis. This study establishes a new link between the nuclear envelope and the regulation of chromosome separation, which contributes to ensure proper chromosome segregation and subsequent genome stability.

## MATERIALS AND METHODS

### Cell cultures

HeLa cells stably expressing H2B-eGFP-Rad21^128-268^-mCherry (separase sensor) were kindly provided by Prof. Dr. Olaf Stemmann, and generated and grown as described in (Henschke et al. 2019). Normal primary adult fibroblasts (GM08399), normal embryonic diploid fibroblasts (WI-38) and immortalized SV40-fibroblasts (GM0639) were purchased from Coriell Cell Repositories, and cultured in Dulbecco’s modified Eagle’s medium (DMEM) medium supplemented with 10% fetal bovine serum (Gibco), 2 mM glutamine, 200 U/ml penicillin (Sigma) and 200 mg/ml streptomycin (Sigma) under 37°C with 5% CO2. Lack of senescence in cell cultures of WI-38 (PD15 to 22) and GM08399 (PD<15) was assessed by SA- β-galactosidase assays at day 0 of each experiment (data not shown).

### Plasmid constructs

Ds-Red lamin B1–expressing vector is described in (Delbarre et al. 2006). Untagged lamin B1 full length was purchased from Origene (#SC 116661). FarRed lamin B1-expression vector was obtained by replacing the eGFP coding sequence from a eGFP lamin B1-expressing vector, by a miRFP670 coding sequence (from pmiRFP650, addgene plasmid #79987) using sequence and ligation independent cloning (SLIC) technique. HA-ESPL1 expressing vector was purchased from Genscript (clone ID: J53779 into pcDNA3.1(+)-N-HA). DNA sequencing was performed to verify each construct.

### Transfections

24 h prior to transfection, SV40-fibroblasts were plated in 6-well plates at the density of 1,75×10^5^/well for most of the experiments or at the density of 1×10^6^ cells/ 100 mm petri dishes for co-immunoprecipitation. Single transient transfections in these cells was achieved using JetPEI or JetPRIME (Polyplus) transfection reagent, according to manufacturer’s instruction recommendations, with an empty plasmid as control (CTRL), a or a pCMV6 plasmid containing the WT human lamin B1 cDNA (LMNB1) or other lamin B1 constructs (DsRed-LMNB1, FarRed-LMNB1). For rescue experiments with HA-ESPL1 vectors, SV40-cells were transfected either with empty CTRL vector, DsRed-LMNB1 vector or a pcDNA3.1(+)-N-HA plasmid containing the WT human separase cDNA (HA-ESPL1) or co-transfected with both LMNB1 and HA-ESPL1 vectors with equal number of moles (2.5 × 10^−13^ mol of DNA/well). For single transfection conditions, the amounts of plasmids were adjusted with CTRL vector to transfect equal moles of DNA in a ratio 1:1 for each condition. For transient transfections on WI-38, cells (5×10^5^/well in 6-well plates) were transfected with 2 μg of DNA by nucleofection using Amaxa device (Lonza) following manufacturer’s instructions. For SiRNA interference, cells were transfected with Interferin reagent (Polyplus) using 10 nM smart-pool ON-TARGETplus siRNAs designed against lamin B1 (Dharmacon) or a scrambled siRNA as control (Eurogenetec) per well in a 6-well plate. Transfection efficiencies were verified by western blotting and immunofluorescence staining using specific antibodies, after 48 h for most experiments.

### Immunofluorescence

Cells grown on coverslips. 48h or 72h after transfection, were fixed in 4% PFA or ice-cold 100% methanol (10 min). After a blocking step in 2% BSA-0.05% Tween-20 in PBS, cells were incubated for 1-2 h at room temperature with the indicated primary antibodies: rabbit anti-lamin B1 (ab16048, Abcam), mouse anti-separase (6h6, Novus Biologicals), mouse anti centrobin (ab70448, Abcam) or rabbit anti HA-tag (ab9110, Abcam). After washes in PBS, cells were incubated 0.75-1 h with the conjugated secondary antibodies (alexa fluor-488, 594 or 647, Life Technologies), followed by washes in PBS and counterstaining with DAPI. Slides were then mounted with fluoromount mounting medium (Southern Biotech). Images were acquired with an epifluorescence microscope leica DM5500B or a Leica spinning disk using a 40×-dry or a 60x-oil objective and analyzed with ImageJ and cell profiler software.

### Proximity ligation assay (PLA)

Cells grown on coverslips were fixed with cold methanol for 10 min, blocked with 2%BSA-0.05%Tween-20 in PBS and incubated with the following primary antibodies couples: mouse anti-separase (6h6, Novus Biologicals) and rabbit anti-lamin B1 (ab16048, Abcam). Next, PLA was performed with the Duolink *in situ* PLA probes (ant-mlouse and anti-rabbit probes, Sigma-Aldrich) and Duolink Red detection Kit (Sigma-Aldrich) following the recommended manufacturer’s protocol. For some experiments, when indicated in figure legends, PLA was coupled with immunostaining of lamin B1. Digital images were acquired with the Leica SP8 confocal microscope using a 60×-oil objective lens or Leica spining using a 40×-dry. Images were processed with ImageJ and cell profiler software.

### Cytogenetic analysis

Metaphase spreads were performed as described previously (Pennarun et al. 2021). Cells (72 h post-transfection for GM0639 and an additional 72h for WI38) were arrest in colcemid 0.1 μg/ml (Sigma) for 2 h at 37°C. Then, cells were collected and incubated in prewarmed hypotonic solution containing 0.0375M KCl and 1:12 human serum (Lonza) for 20 min at 37°C. After fixation in 3:1 (v/v) ethanol/acetic acid, cells were dropped onto ice-cold slides and air-dried overnight at room temperature. Metaphase spreads were stained with 4% Giemsa stain (Sigma) or DAPI and captured using bright-field or fluorescent microscopy (DM5500B, Leica), respectively, with a 60×-oil objective lens or alternatively.

### Western-blot

To extract proteins, cells were homogenized in ice-cold SDS buffer (10 mM Tris (pH 7.5), 1% SDS) supplemented with 1X complete protease inhibitor cocktail (Roche) and phosphatase inhibitors cocktail 2 and 3 (Sigma) one ice for 1 h, as described previously (Pennarun et al. 2021). 30 to 50 μg of protein extracts were resolved by SDS-polyacrylamide gel electrophoresis using 4–12% NuPAGE Bis-Tris gradient gel or 3–8% Tris-acetate gel and run (Invitrogen) and MOPS or Tris-acetate running buffer (Invitrogen) and further blotted onto nitrocellulose membrane (Amersham). After a blocking step with 5% of non-fat milk, membranes were probed with the indicated primary antibodies: mouse anti-separase (6h6, Novus Biologicals), rabbit anti-lamin B1 (ab16048, Abcam), rabbit anti-phospho-histone H3 (Ser10) (06-570, Upstate). Rabbit anti-ß-actin (2066, Sigma) or mouse anti-Vinculin (ab18058, Abcam) antibodies were used as loading controls. Primary antibodies were detected using anti-mouse or -rabbit secondary fluorescent antibodies coupled to IRDye700 or IRDye800 (Diagomics). Fluorescent signals were detected using Odyssey Imaging system (LI-COR) or iBright system (Thermo Fisher) and protein levels were quantified by densitometric analysis using Image Studio or iBright Analysis Software software. Levels of protein of interest were normalized with respect to loading control.

### Co-immunoprecipitation

SV40-fibroblasts cells were lyzsed in ice-cold lysis buffer containing 50 mM Tris–HCl (pH 7.5), 150 mM NaCl, 5 mM EDTA, 0.5% NP40 supplemented with 1X protease and phosphatase inhibitor cocktails (Roche) for one hour on ice. After centrifugation (13 200 rpm/10 min at 4°C), supernatants containing protein extracts were pre-treated with the Benzonase nuclease (0.5 U/μl, E1014, Sigma) supplemented with 10 mM MgCl2 for 1 h at room temperature (RT) and a fraction of extracts was kept as input. 5 μg of rabbit antibodies specific to lamin B1 (ab16048, Abcam) or mouse anti-ESPL1 (6H6, Novus Biologicals) were coupled with Dynabeads protein G (10007D, Life Technologies) for 1 h at RT. Mouse or rabbit preimmune IgG (5 µg, Santa Cruz) were used as negative controls. After 3 washes in PBS- 0.05% Tween-20, coupled beads were incubated with 0.5–1 mg of protein extracts for 1.5 h at RT, and washed 4 times in Dynabead Washing Buffer (Life technologies). Then, after elution in Laemmli buffer (2X)/ 4% ß-mercaptoethanol, protein extracts and inputs (60 µg) were resolved by Western blotting as described above using indicated primary antibodies.

### Cell synchronization

For synchronization in mitosis, cells were treated with 2mM thymidine (Sigma-Aldrich) for 16h, then washed in fresh medium and released for 9h, and treated a second time with 2mM thymidine for 16h. After the release from the second block, cells entered mitosis in the next 8 to 10h. When coupled with transfection, cells synchronization was started 6 hours after transfection, as described below. Synchronization efficiency was assayed by analyzing the enrichment of pH3^S10^ by western blot.

### Time lapse microscopy

For time-lapse microscopy, cells were seeded in 8-well chambered coverslip (Ibidi GmbH, Germany) at the density of 35000 cells per wells. Cells were then transfected and synchronized as described above. At the time of acquisition, medium was changed for a red phenol-free medium (Gibo). Acquisitions were then started 6 h after the release of the second thymidine block. Images were acquired every 2.5 min during 16 h, with 150 ms exposure times, with a Leica spining disk, 20X dry objective. During the time-lapse, cells were kept at a constant temperature of 37°C, constant O_2_ and CO_2_ concentration (20% and 5%, respectively). For data analysis, we used Cell Profiler software to track mitosis over time, using the mCherry signal to segment individual cells. Single mitosis were then aligned on the frame corresponding to the onset of their nuclear envelope breakdown, determined by the first frame with detectable chromosome condensation. The ratio between mCherry and GFP signal in time-lapse microscopy provides a precise measure of the evolution of ESPL1 enzymatic activity over time, this value is termed Rcut = 1- (GFP/mCherry) as described in Shindo et al. 2012. Rcut values were then subjected to min-max normalization to fit between 0 and 1. For quantification of the time between the nuclear breakdown (NEB) and anaphase onset, NEB onset was determined for each cell analyzed in using the first frame with visible chromosome condensation and nuclear collapse on the mCherry signal; anaphase onset was determined using the first frame with a clear separation of the two sets of chromosomes

## Acknowledgments

We thank all the members from P.B. lab and G.C lab for advice and support. We acknowledge the core facilities of the iRCM institute. We are grateful to Pr Olaf Stemmann and Susanne Hellmuth for providing us the HeLa cell line with the separase substrat. This work was supported by the “Institut National du Cancer” (INCa_18519), Emergence University Paris-City grant, Institut National de la Santé et de la Recherche Médicale (INSERM) house funding for P.B lab. This work has been also supported by a grant of AFM-Téléthon obtained by G.C. and P.B and grants from Institute Pasteur of Rome (“A. Tramontano 2022”) and Italy Ministry of University and Research (PRIN, N 202227SYBW) to G.C, a grant from Italy Ministry of University and Research (PRIN, N 2022KKMTL9) to F.C. JP was supported by a Ph.D Fellowship from the French Ministry of Higher Education Research and Innovation (MESRI) and a 4rd year PhD grant from the “Association pour la Recherche contre le Cancer” (ARC). The illustrations (Fig. 6A and Fig. 8) were created with BioRender.com.

## Author Contributions

GP, PB, JP: conceived the study; FC, GC: conceptualized the original idea of the separase-lamin link; GP, JP: designed, performed most of the experiments and data analysis; DB: created and verified most of the plasmid constructs used in this study; GP and PB: supervised the study; JP and GP: wrote the original draft of the manuscript; GP, PB, FC, GC: re-viewed and edited the manuscript.

